# *In vivo* brain temperature measurement using quantum dot imaging temperature sensors

**DOI:** 10.64898/2026.06.11.731502

**Authors:** Mariko Handa, Makoto Tozawa, Fuyu Miyaji, Shota Yamada, Masaki Yoshioka, Manami Takahashi, Yasuyuki Ueda, Shigetatsu Tsugimoto, Kazutaka Akiyoshi, Ayaka Takada, Shuri Takemoto, Chihiro Ito, Taisuke Shimada, Yuki Watakabe, Hirokazu Ishii, Motosuke Tsutsumi, Tomomi Nemoto, Jeff Kershaw, Tatsuya Kameyama, Masazumi Fujiwara, Yoshinobu Baba, Masakazu Agetsuma, Tsukasa Torimoto, Hiroyuki Takuwa, Hiroshi Yukawa

## Abstract

Temperature regulation in the brain is essential for maintaining neuronal function and preventing thermally induced damage. Here, we report the development and *in vivo* application of quantum dots (QDs) to high-resolution thermometry in the mouse brain using two-photon excitation microscopy. These QDs, via the red-to-green photoluminescence (PL) intensity ratios, enabled stable temperature measurements in both normal and chronically hypo-perfused cerebral tissue. Our findings show that localized neuronal activity leads to transient heat generation, which is rapidly dissipated by cerebrovascular responses. In a chronic hypoperfusion model, impaired vascular function resulted in exaggerated and prolonged brain temperature elevations. This thermometry system provides unprecedented insight into the mechanisms of cerebral thermoregulation and highlights the importance of vascular cooling in protecting the brain from heat-induced stress, particularly in pathological conditions such as stroke.

## Introduction

Temperature-sensitive fluorescent tracers can be miniaturized to the nanoscale and measured non-invasively via microscopy, making them particularly effective for detecting micro-regional temperature dynamics. To date, various nanoscale temperature probes such as fluorescent proteins^1^, nanodiamonds^2^ and polymer nanoparticles^3^ have been developed. However, due to limited photoluminescence (PL) intensity and biological permeability, nanoscale temperature sensors of this sort have mainly been applied to measuring the temperature of cultured cells or the surface of the body.

Quantum dots (QDs) are semiconductor nanoparticles with tunable emission wavelengths, and temperature-sensitive variants have been developed and applied in several *in vivo* thermometric studies^4–7^. Conventional temperature-sensing QDs rely on the correlation between PL intensity and temperature, which unfortunately makes them susceptible to optical artifacts such as absorption and scattering by biological tissues. This issue is particularly critical in detailed *intra-vital* imaging using two-photon microscopy, where the optical environment can change over time due to dural thickening beneath the cranial window, potentially compromising the reliability of temperature measurements.

The brain constitutes a large-scale neural circuit in which a vast number of neurons actively produce heat through metabolic activity. Although it represents only about 2% of body weight, the brain consumes roughly 20% of total oxygen intake^8–10^ and this intense oxidative metabolism leads to substantial endogenous heat production^9,11^. To preserve cerebral homeostasis, the heat must be effectively dissipated^11^. There is evidence that cerebral blood flow (CBF), accounting for 15–20% of cardiac output^12,13^, plays a key role in absorbing excess parenchymal heat and transporting it away. Specifically, the inflow temperature of the carotid artery is known to be ∼0.3°C lower than that of venous outflow under resting conditions^14–17^, suggesting that the highly vascularized cerebral network has an active heat exchange function.

In contrast to other organs, the brain exhibits a unique spatiotemporally dynamic pattern of temperature change, the source of which is neuronal activity^18^. However, the mechanisms that regulate this dynamic thermal activity remain poorly understood. Localized increases in CBF associated with neuronal activation, known as neurovascular coupling^19,20^, have long been recognized. These increases arise from both direct neuronal signals and indirect pathways mediated by astrocytes^21^, leading to vasodilation of nearby arterioles and capillaries. This suggests that the brain uses its vascular architecture as a highly sophisticated “localized water-cooling” system to manage transient thermal hotspots generated by neuronal firing, but no direct proof of that hypothesis exists. One way to directly validate this hypothesis would be to develop microscale in vivo sensors capable of real-time temperature measurement under physiological conditions where neuronal and vascular functions are preserved. Furthermore, to measure temperature within the brain parenchyma, the sensor must be compatible with two-photon excitation. Compatibility with two-photon microscopy would allow volumetric, four-dimensional temperature imaging of deep brain parenchyma—an area previously difficult to access. In contrast to state-of-the-art nanodiamond-based quantum sensors, which offer temporal resolution only to the scale of seconds and require complex microwave synchronization and data analysis, the quantum dots developed in this study achieve millisecond-level temporal resolution and micron-level spatial resolution, while being compatible with standard two-photon microscopy setups. This makes them a highly versatile and practical tool for deep-tissue thermal imaging.

In this study, we have successfully developed low-toxicity quantum dots (AgInGaSe@ZnGaS: AIGS@GZS QDs), which simultaneously exhibit both band-edge and defect-site emission peaks, with a peak intensity ratio that changes linearly with the surrounding environmental temperature. Because the technique relies on the temperature-dependent PL intensity ratio rather than absolute intensity, it reduces the potential influence of tissue-induced optical artifacts that trouble conventional methods, thereby enabling stable *in vivo* temperature measurements.

The feasibility of temperature measurement in live mice using AIGS QDs was first confirmed with an *in vivo* PL imaging system. AIGS QDs were then applied to observe cortical temperature dynamics in live mice using two-photon microscopy. In response to sensory stimulation of the whisker pad, we observed changes in somatosensory cortical neuronal activity, blood flow, and temperature. The same methodology was also applied to a mouse model of chronic cerebral hypoperfusion, where vascular function is impaired, to evaluate cerebrovascular contributions to thermal regulation. The results provide direct evidence for a vascular-mediated cooling mechanism in the brain and suggest that disruption to its function may exacerbate heat-related injury in stroke. This high-spatiotemporal-resolution thermometry system offers a powerful tool for investigating brain temperature regulation and might also be used to study thermal physiology in other organs.

## Materials and Methods

### 2.1. Materials

Silver (I) acetate (Ag(OAc)), zinc (II) stearate, and 1-dodecanethiol (DDT) were purchased from FUJIFILM Wako Pure Chemical Corp. Indium (III) acetylacetonate (In(acac)^3^), gallium (III) acetylacetonate (Ga(acac)^3^), oleylamine (OLA), and tetramethylammonium hydroxide solution (TMAOH, 25wt% in methanol) were purchased from Sigma-Aldrich. Thiourea and sulfur powder were purchased from Kishida Chemical Co., Ltd. and 3-mercaptpropionic acid (MPA) was obtained from Tokyo Chemical Industry Co., Ltd.

### 2.2. Animals

Female BALB/c mice (20-30 g; 6 weeks of age; Japan SLC Inc., Hamamatsu, Japan) were used for subcutaneous temperature measurement of mouse. Male C57BL/6J mice (20-30 g; 7-11 weeks of age; Japan SLC Inc., Hamamatsu, Japan) were used for two-photon microscopic immunohistochemistry examinations. Male double-transgenic mice (20-30 g; aged >7 weeks) expressing GCaMP6s and CaMKIIα-Cre were used for *in vivo* calcium imaging. The mice were obtained by mating GCaMP6s transgenic mice (B6J.Cg-Gt(ROSA)26Sor<tm96(CAG-GCaMP6s)Hze>/MwarJ, No. 028866, The Jackson Laboratory, Sacramento, California) and CaMKIIα-Cre transgenic mice (B6.Cg-Tg(Camk2a-cre) T9-1Stl/J, No. 005359, The Jackson Laboratory, Sacramento, California). The animals were housed with food and water provided ad libitum in cages maintained at 25°C under a 12-hour light/dark cycle. All experiments were conducted in accordance with institutional guidelines for the humane care and use of laboratory animals and were approved by the Institutional Committee for Animal Experimentation of the National Institutes for Quantum Science and Technology.

### 2.3. Synthesis of Ag–In–Ga–S core/ Ga– Zn–S shell-structured QDs (AIGS@GZS QDs)

QDs composed of Ag–In–Ga–S (AIGS) solid solution were synthesized by a previously reported method with slight modifications^22^.The precursor metal ion ratios were Ag^+^/(Ag^+^ + In^3+^ + Ga^3+^) = 0.40 and In^3+^/(In^3+^ + Ga^3+^) = 0.60, while the charge ratio of sulfide ions to total metal ions was fixed at 0.90. A mixture of Ag(OAc) (0.080 mmol), In(acac)_3_ (0.075 mmol), Ga(acac)_3_ (0.050 mmol), and elemental sulfur (0.23 mmol) was put in a test tube, followed by the addition of OLA (2.75 cm^3^) and DDT (0.25 cm^3^). The solution was heated at 300℃ for 10 min under vigorous stirring in an N_2_ atmosphere. After cooling for 8 min at room temperature, the resulting suspension was centrifuged to remove aggregated large particles. The target AIGS QDs were precipitated from the supernatant by adding methanol as a non-solvent, followed by centrifugation. The precipitate was washed with ethanol and finally dissolved in chloroform.

The thus-obtained AIGS QDs were used as a core and their surface was coated with a Ga–Zn–S shell layer based on a previous method with slight modifications^23^. Portions of the AIGS QDs (1.0 × 10^−5^ mmol), zinc stearate (0.32 mmol), Ga(acac)_3_ (0.16 mmol), and thiourea (0.56 mmol) were added to 3.0 cm^3^ of OLA. The mixture was vigorously stirred at 150 ℃ for 10 min in an N_2_ atmosphere and then immediately heated at 250 ℃ for 30 min. After cooling to room temperature, the reaction solution was centrifuged to precipitate Ga–Zn–S-coated AIGS QDs, which were subsequently washed with methanol and ethanol. The resulting QDs were further treated with ZnS to form a Ga–Zn–S shell layer with a Zn-rich surface. A portion of the QDs (5.0 × 10^−6^ mmol) was put in OLA (3.0 cm^3^) along with zinc stearate (0.16 mmol) and thiourea (0.16 mmol). The mixture was subjected to a two-step heating process at 150 ℃ and 250 ℃, following a procedure similar to that used for the Ga–Zn–S shell coating described above. The resulting QDs, having a AIGS core/Zn-rich Ga–Zn–S shell structure, AIGS@GZS, were then isolated and purified before being dissolved in 3.0 cm^3^ of chloroform.

The QDs synthesized in OLA were hydrophobic due to the adsorption of an OLA layer on their surface. Thus, to enable uniform dispersion in aqueous solution, the initial surface ligands were exchanged with MPA using a modified version of a procedure reported in a previous paper^24^. After two-fold dilution of the QD chloroform solution with chloroform, 1.0 cm^3^ of the resulting solution was added to a mixture of TMAOH methanol solution (0.73 cm^3^), ethanol (0.27 cm^3^), and MPA (0.10 cm^3^). The resulting suspension was heated at 70℃ under stirring in an N_2_ atmosphere for 3 h. After cooling to room temperature, the suspension was mixed with acetone, followed by centrifugation. The resulting wet precipitates of MPA-modified AIGS@GZS QDs were washed several times with acetone and then dispersed in pure water. To remove aggregated particles, the solution was filtered through a membrane filter (pore size: 0.20 μm) before use in further experiments (Figure 1a).

**Figure 1:**
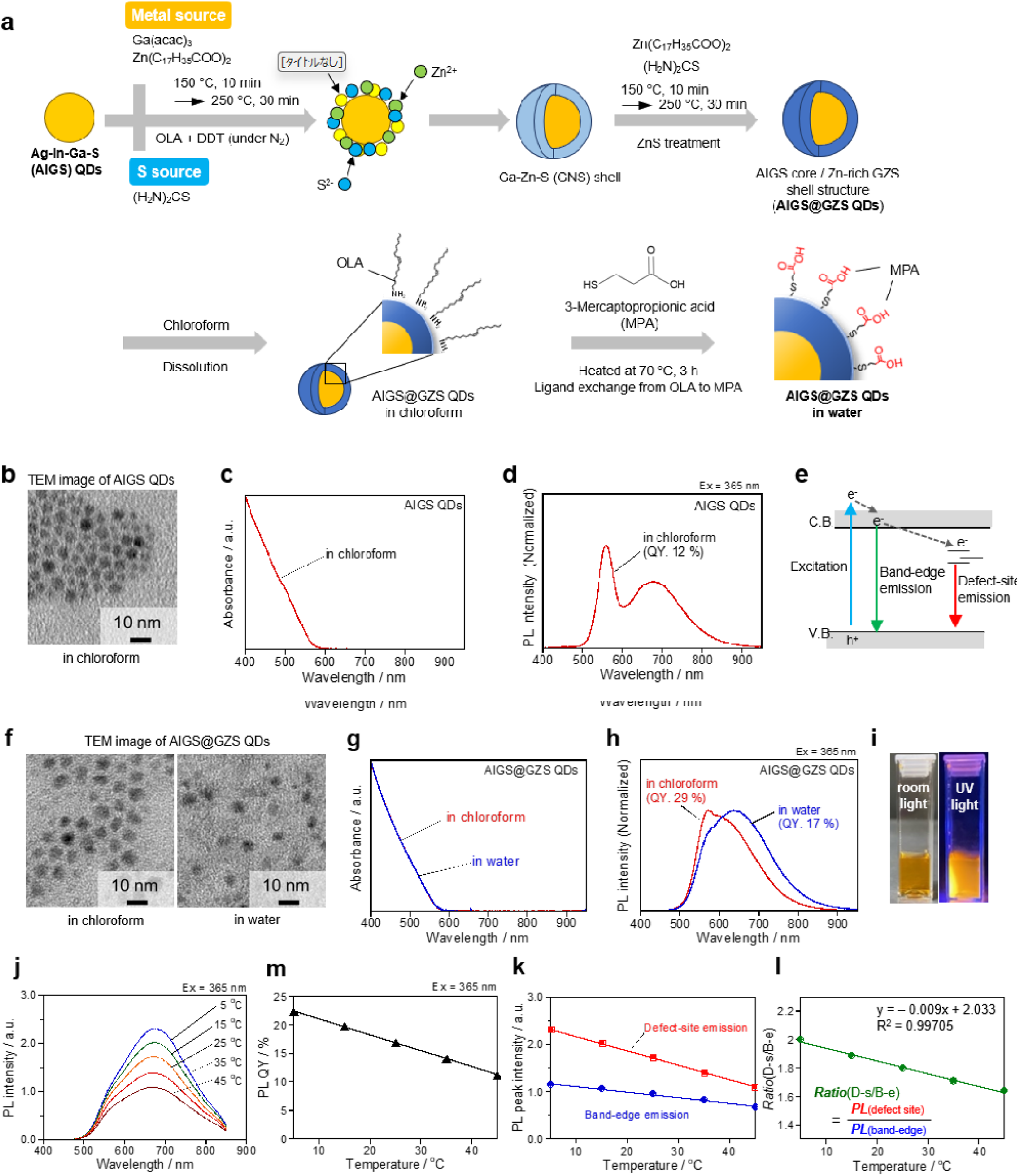
(a) Schematic illustration of the preparation of water-dispersible AIGS@GZS QDs. (b) TEM image of AIGS QDs. (c,d) Absorption (c) and PL spectra (d) of AGIGS QDs in chloroform. (e) Schematic illustration of the mechanisms of band-edge and defect-site emission. (f,g,h) TEM images (f), absorption spectra (g), and PL spectra (h) of AIGS@GZS QDs. The QDs modified with OLA and MPA were uniformly dispersed in chloroform and water, respectively. (i) Photographs of an aqueous solution containing MPA-modified AIGS@GZS QDs under ambient light and UV light. (j,m) PL spectra (j) and PL QYs (m) of MPA-modified AIGS@GZS QDs in water at various temperatures. (k,l) Temperature dependence of (k) the intensities of band-edge and defect-site emission in panel j and (l) the corresponding Ratio(D-s/B-e) values.

### 2.4. Optical characterization

Absorption spectra were measured using an Agilent 8453A diode array spectrophotometer. PL spectra at room temperature were recorded using a photonic multichannel analyzer (Hamamatsu, PMA-12, C10027-02) with 365 nm excitation light. The PL quantum yield (QY) at room temperature was evaluated using an absolute PL QY measurement system (Hamamatsu, C9920-03) with 365 nm excitation. Temperature-dependent PL spectra were acquired with a spectrofluorometer (HORIBA, FluoroMax-4) with excitement at 365 nm. Measurements were conducted in the temperature range of 5-45 °C using a temperature-controlled cell holder connected to a circulating chiller (EYELA, NCB-1210). The relative PL QY of the QDs at each temperature was estimated from the integrated PL spectral area, using the absolute PL QY of the QDs measured at room temperature (25 °C) as the reference.

### 2.5. Transmission electron microscopy (TEM)

To analyse the size distribution of the QDs, wide-area TEM images were acquired using a transmission electron microscope (TEM, Hitachi, H-7650) operating at an accelerating voltage of 100 kV. TEM samples were prepared by dropping a QD solution on a copper TEM grid covered with an amorphous carbon overlayer (Okenshoji Co., Ltd., ELS-C10 STEM Cu100P grid). The size distribution of the QDs was evaluated using image processing software (ImageJ).

### 2.6. Subcutaneous temperature measurement of mouse

The temperature of AIGS@GZS QDs dissolved in 50 μL of distilled water in a microtube was varied from 10.5°C to 49.8°C using a temperature controller consisting of a Peltier element, and PL images and temperature were measured at each point (Figure 2b, d). In addition, AIGS@GZS QDs dissolved in 50 μL of distilled water were injected subcutaneously into a euthanized BALB/c nude mouse (Figure 2c). The temperature of the mouse varied from 10.5°C to 49.8°C, and PL images and temperature were measured at each point (Figure 2f). The temperature was measured using an infrared camera (FLIR C2, Teledyne FLIR LLC, USA), while the PL imaging was performed using an IVIS Lumina K Series III system (PerkinElmer Inc, Waltham, MA, USA; excitation band filter: 430–450 nm, emission band filters: 550-590 nm, 690-730 nm) (Figure 2a). The PL images were analyzed using ImageJ to obtain the PL intensity at each temperature. The PL intensity ratio was calculated by dividing the band-edge emission intensity (550–590 nm) into the defect-site emission intensity (690–730 nm). The temperature responsiveness was evaluated based on the correlation between the observed PL intensity ratio and the measured temperature, and a calibration curve was constructed (Figure 2e, g).

**Figure 2:**
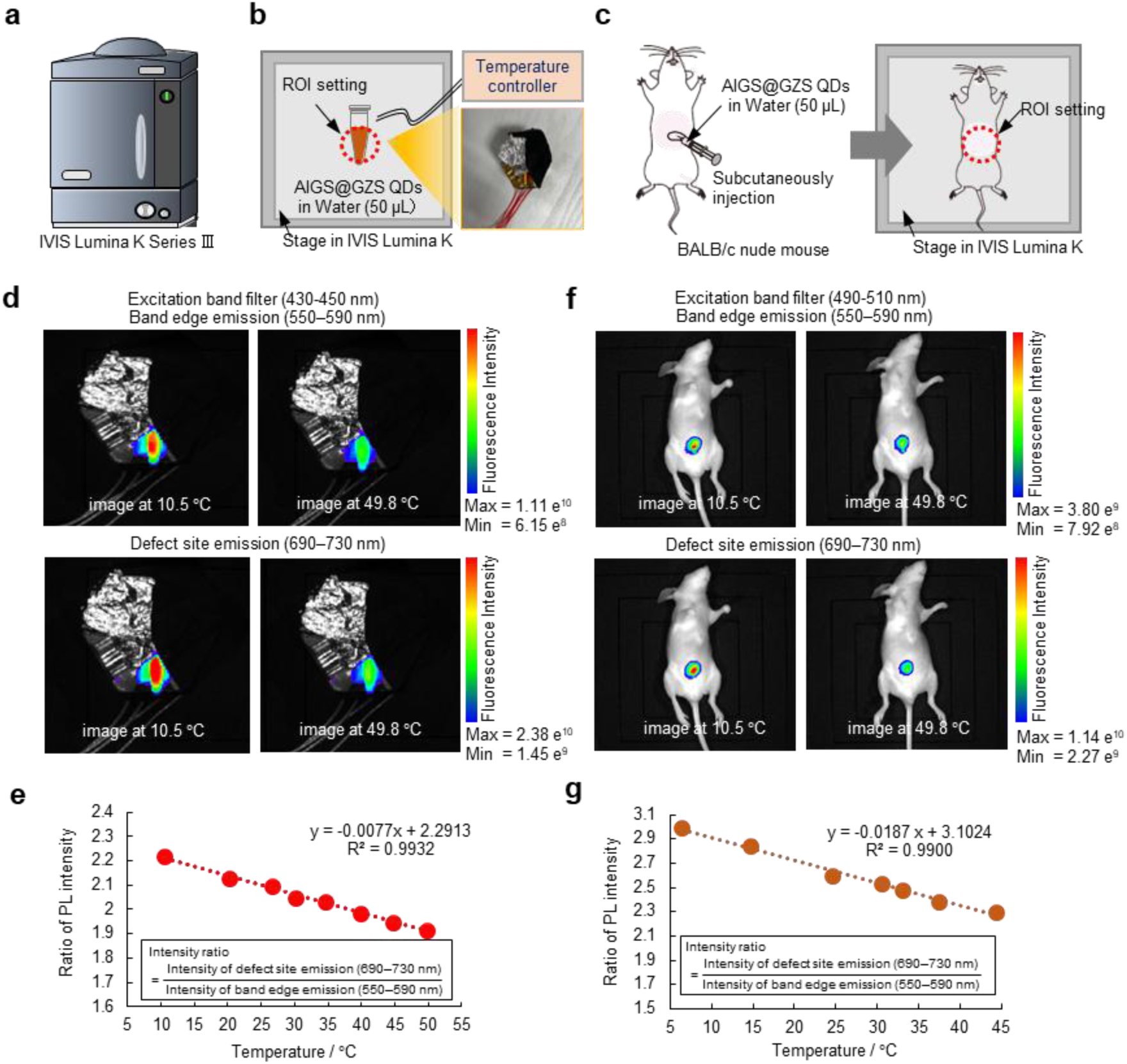
(a) Schematic diagram of the PL imaging system. (b) Schematic diagram of PL imaging of AIGS@GZS QDs dissolved in 50 μL of distilled water in a microtube. (c) Schematic diagram of PL imaging of a euthanized mouse subcutaneously injected with AIGS@GZS QDs. (d) PL images of AIGS@GZS QDs in a microtube at 10.5°C (left) and 49.8°C (right), acquired using an excitation band filter: 430–450 nm and emission band filter: 550-590 nm (upper panel) and 690-730 nm (lower panel). (e) Correlation plot between temperature and the PL intensity ratio of AIGS@GZS QDs in a microtube (R² = 0.9932). (f) PL images of a euthanized mouse subcutaneously injected with AIGS@GZS QDs at 10.5°C (left) and 49.8°C (right), acquired using an excitation band filter: 430–450 nm and emission band filter: 550-590 nm (upper panel) and 690-730 nm (lower panel). (g) Correlation plot between temperature and the PL intensity ratio of a euthanized mouse subcutaneously injected with AIGS@GZS QDs (R² = 0.9900).

Furthermore, AIGS@GZS QDs in 50 μL of PBS were injected subcutaneously into the abdomen of live BALB/c nude mice. Temperature and PL images were obtained using the same method as described in the previous paragraph under three conditions: at room temperature, after cooling the abdomen with ice for 5 min., and after heating with a heat pack for 10 min. The temperature was measured by calculating the PL intensity ratio and using the calibration curve (Figure 2g). The result was then compared with the temperature measured using a near-infrared camera (Figure 3a-c).

**Figure 3:**
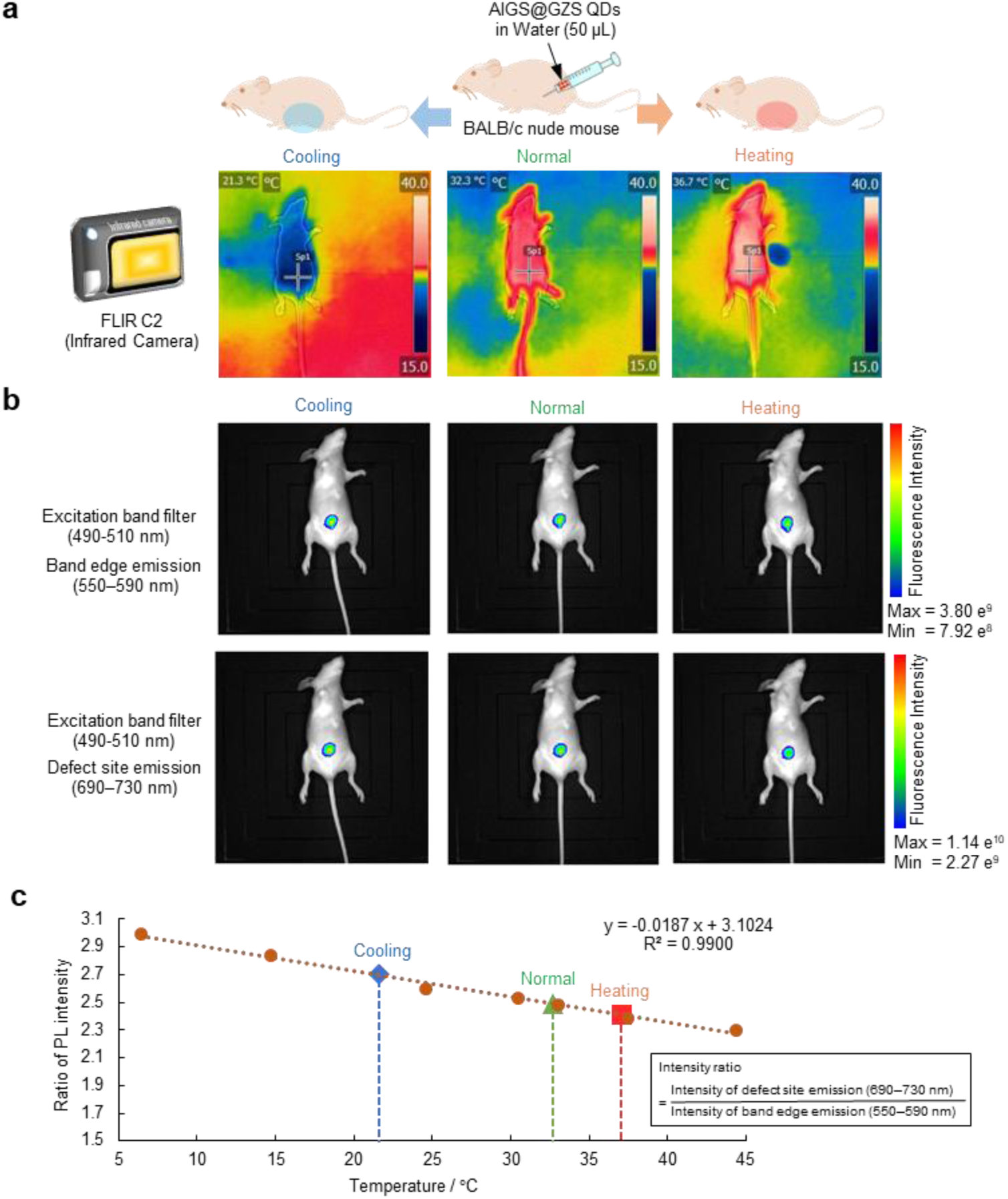
(a) Schematic diagram of the *in vivo* temperature sensing ability of AIGS@GZS QDs in a live mouse, and thermographic images of a mouse at room temperature (center: 32.3°C), after cooling (left:21.3°C) and after heating (right:36.7°C). (b) PL images of a live mouse subcutaneously injected with AIGS@GZS QDs at room temperature (center), after cooling (left) and after heating (right), acquired using an excitation band filter: 430–450 nm and emission band filter: 550-590 nm (upper panels) and 690-730 nm (lower panels). (c) Temperature derived from the PL intensity ratio using the calibration curve.

### 2.7. NIR laser excitation of AIGS@GZS QDs

To evaluate the wavelength response to NIR laser excitation, a 5 μL sample of a solution containing 50 μM of AIGS@GZS QDs was placed between a slide glass (S1112, Matsunami) and a cover glass (18 × 18mm No.1S, Matsunami) using a Frame-Seal^TM^ Incubation Chamber (SLF0201, BioRad). The sample solution was excited using a femtosecond-pulsed, mode-locked Ti:sapphire laser (Mai Tai DeepSee eHP, Spectra-Physics), which was directed onto the sample via an upright microscope (A1R-MP^+^; Nikon) equipped with a water immersion objective lens (CFI75 LWD 16X W, 0.8 NA, Nikon). The laser power was adjusted to be 15 mW at each measured wavelength (700–1040 nm, 10 nm step), by measuring the intensity of the light after passing through the objective lens using a microscope power sensor head (S170C, Thorlabs). The emission at each excitation wavelength was detected with or without optical filters: FF01-520/35 (Semrock) for “green”, NDD592 (Nikon) and BSP01-633R (Semrock) for “red”, (Figure 4a-c). Images were acquired at a resolution of 512 × 512 pixels (793 µm × 793 µm field of view). The emission at a depth of 100 µm from the surface of the sample (i.e., the bottom of the cover glass) was used as the representative signal level, while the signal intensity at 100 µm above the cover glass was measured as the background level. Sequential measurements across the wavelength range were automated using a macro function in the NIS-Elements software (Nikon). Images were analyzed with ImageJ to obtain the mean intensity over the whole image at each wavelength (Figure 4a-c).

**Figure 4:**
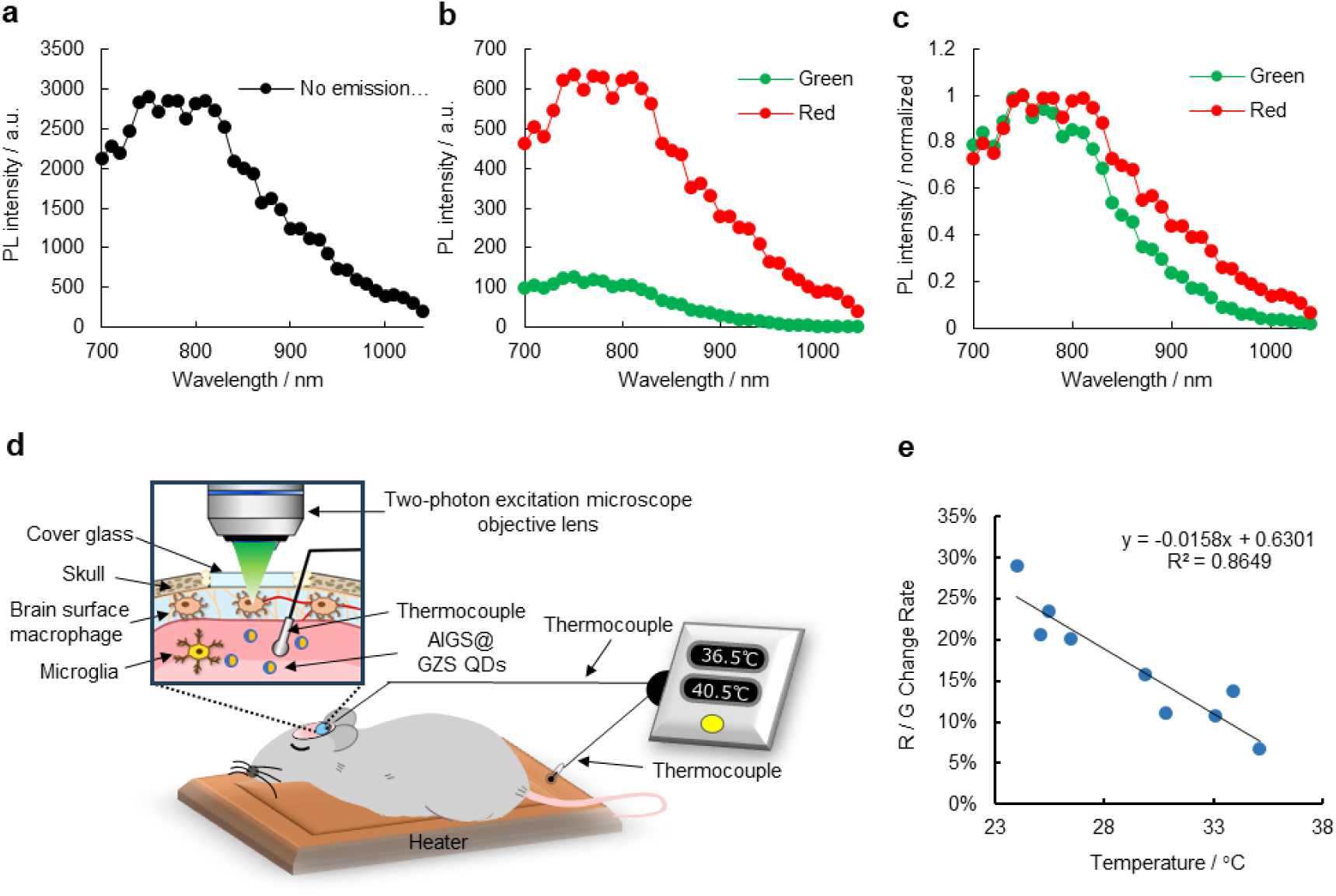
(a–c) Spectral profiles of AIGS@GZS under NIR laser excitation. Absolute (a,b) or normalized (c) emission levels, measured without (a) or with (b,c) emission filters, are shown. (d) Schematic diagram of the calibration of AIGS@GZS QDs using postmortem brains. (e) Graph comparing the temperature measured using a thermocouple probe inserted into the brain parenchyma of a postmortem mouse with the rate of change in PL intensity of AIGS@GZS QDs in the brain parenchyma. QD PL was obtained when the measurement value measured using the thermocouple probe was between 24℃ and 35℃. R² = 0.8649.

### 2.8. Temperature measurement of QD phantom using a two-photon excitation microscope

The following procedures were performed to prepare a 1.5% agarose gel QD phantom Agarose (Agarose S, Nippon Gene Co., Ltd., Tokyo, Japan) and QDs were added to physiological saline at a 200-fold dilution, mixed thoroughly, and poured into a 35 mm glass-bottom dish. The dish was then left undisturbed overnight in a refrigerator (4°C). A temperature probe (thermocouple) was inserted into the phantom, and a coverslip was placed on top. The phantom was then placed on a temperature control device (Thermo Plate III, Tokai Hit, Shizuoka, Japan) and imaged using a two-photon excitation laser microscope (Thorlabs, Newton, New Jersey, USA). All measurements using the two-photon excitation laser microscope were performed with a × 25 water immersion objective lens (NA = 1.0, Olympus). Excitation was performed using a 920 nm laser, and defect PL emission (red) and band-edge PL emission (green) were captured with two separate detectors. The two-photon excitation laser microscope was controlled using the ThorImage^®^ software (Thorlabs) (Figure S3a). The control device gradually increased the temperature, and the PL from the QDs was acquired at each setting. The temperature measured by the thermocouple in the phantom varied from 25°C to 40°C. The ratio of the red-to-green PL intensities was calculated and plotted against the temperature of the thermocouple probe to create a calibration curve (Figure S3b).

Next, the QD phantom was placed on a temperature control device (BWT-100, Bio Research Center Co., Ltd., Aichi, Japan) with a thermocouple temperature probe inserted. The thermocouple was feedback-controlled by the temperature control device. The temperature probe measurements fluctuated continuously between 29.9°C and 30.1°C, when the temperature control device was set to 30°C. Under these conditions, the quantum dot phantom was excited with 920 nm light using a two-photon excitation laser microscope, and PL emissions in the red and green channels were captured. The image resolution was 1024 × 1024 pixels, and measurements were performed at a rate of 0.14 seconds per frame for 60 seconds (Figure S3b).

### 2.9. Calibration curve for QD measurement of mouse brain temperature using a two-photon excitation microscope

To obtain a QD PL intensity versus temperature calibration curve, we performed the following experiment. A C57BL/6J mouse was euthanised after the installation of a cranial window. A glass micropipette was used to inject QDs diluted 200-fold in physiological saline into the brain parenchyma at the same time that a thermocouple temperature probe was inserted simultaneously. The mouse was placed on a temperature control device (STXG-WSKMX-SET, Tokai Hit, Shizuoka, Japan), and imaging was conducted using a two-photon excitation laser microscope (Figure 4d). All measurements using the two-photon excitation laser microscope were performed with a ×25 water-immersion objective lens (NA = 1.0). Excitation was performed with a 920 nm laser, and PL from defect-state emission (red) and band-edge emission (green) was separately captured using two separate detectors. The microscope was controlled using the ThorImage software (Thorlabs). The set temperature of the temperature control device was gradually increased, and PL images from the QDs were acquired at each temperature while the brain temperature measured by the thermocouple ranged between 24°C and 35°C. The temperature was then calculated from the ratio of red-to-green PL intensity was then calculated and plotted against the actual temperature measured by the thermocouple to generate a calibration curve (Figure 4e).

### 2.10. Intracerebral injection of AIGS@GZS QDs

To locally inject QDs into the brain parenchyma, a microinjection needle was prepared using a glass capillary (BF100-50-10, Sutter Instrument, California, USA) and a micropipette puller (PC-100, Narishige, Tokyo, Japan). The prepared needle was attached to one end of a tube (TYGON LMT-55, SAINT-GOBAIN, La Défense, France), while a syringe (Terumo Tuberculin Syringe 1 mL, Terumo Corporation, Tokyo, Japan) was attached to the other end. The assembly was mounted onto a three-dimensional hydraulic micromanipulator, and the needle was inserted into the left somatosensory cortex of the mouse brain under isoflurane anesthesia (3% for induction, 1.5–2% for maintenance). QDs or saline (as a control) were injected into the brain parenchyma (1.0–1.5 µL per site, total of 3–4 sites). Three days after injection, the mice were perfusion-fixed, and their brains were extracted. Coronal brain sections (thickness: 70 µm) were prepared using a vibratome (VT1200S, Leica Microsystems, Tokyo, Japan). Immunofluorescence staining was performed using anti-GFAP antibody (1:2500), anti-Iba1 antibody (1:2000), and anti-tubulin β3 antibody (1:2000). The stained sections were imaged using a Zeiss Observer spinning disk confocal microscope equipped with diode lasers (405, 488, 594, and 647 nm) and the Zen acquisition software (Zeiss) (Figure 5a). To analyze the localization of QDs within the brain parenchyma, tiled images of single optical planes were acquired at 10× and 20× magnifications, and the field of view (FOV) was selected using the Zen acquisition software (Figure 5b, c).

**Figure 5:**
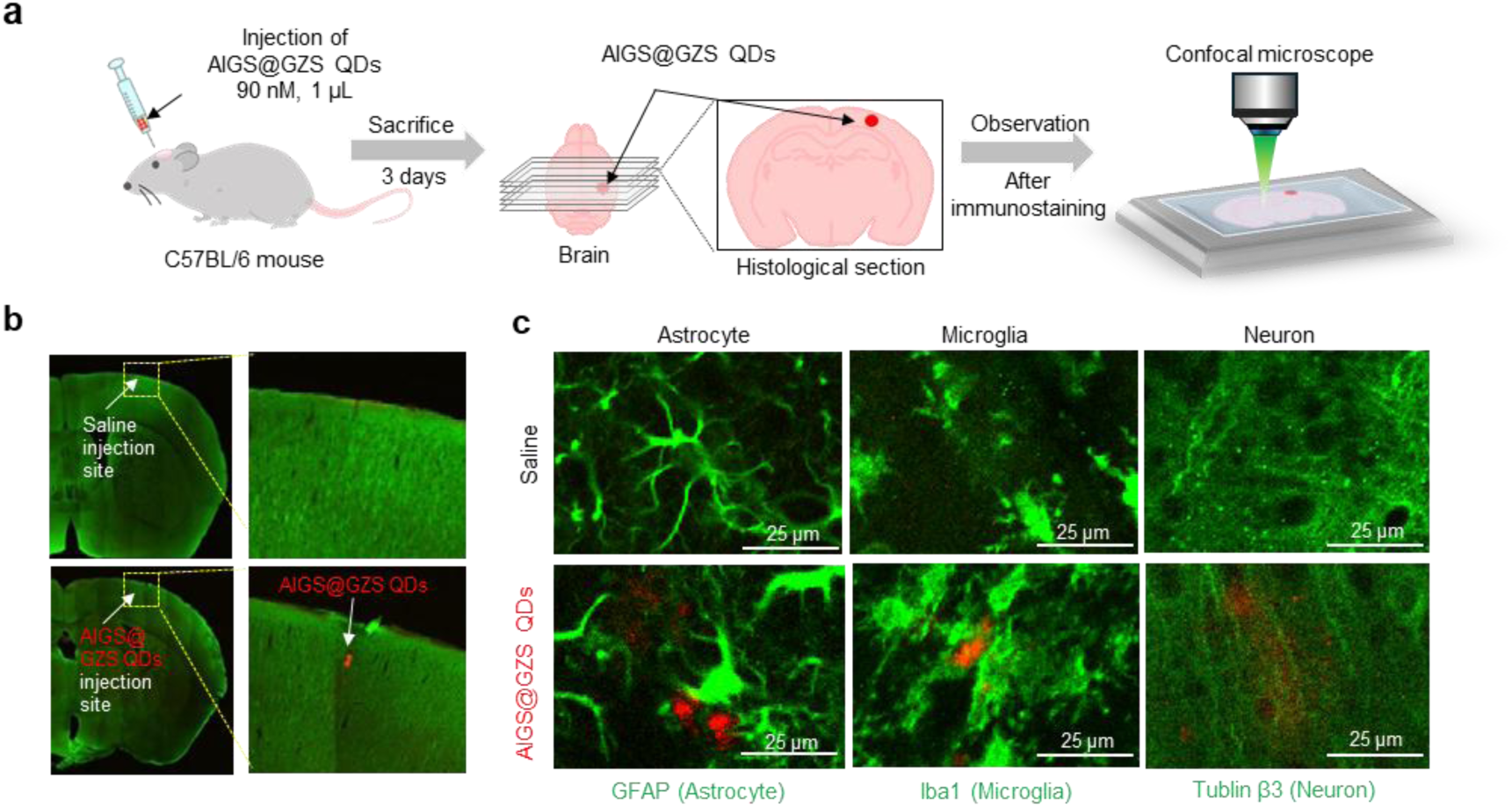
(a) Schematic diagram of the experimental method to investigate the distribution of QDs injected locally into the brain. (b-c) Representative confocal images of tissue sections of the brain parenchyma after injections of saline (upper panel) or AIGS@GZS QDs (red) (lower panel). (b) Left: immunostained image of a brain hemisphere. Right: enlarged view of the injection site. (c) Representative images of fluorescent immunostaining of astrocytes (left), microglia (center), and neurons (left). Scale bar, 25 µm.

### 2.11. Cranial window installation and measurement of temperature in mouse brain parenchyma

During the surgical preparation for *in vivo* temperature measurement, the C57BL/6J mice were anesthetized with a mixture of air, oxygen, and isoflurane (3–5% for induction and 2% for surgery) via a facemask. The animal’s head was fixed in a stereotactic frame. Rectal temperature was monitored, and the animal was heated to maintain body temperature at 36-37°C. The cranial window was attached over the left somatosensory cortex, centered at 1.8 mm caudal and 2.5 mm lateral from the bregma, using dental cement (Luxaflow, DMG, Hamburg, Germany). During this surgery, AIGS@GZS QDs (90 nM, 1 μL) were injected into the barrel cortex using a glass needle. Two custom-made 14 mm long plastic bars were affixed to the skull so that the animal’s head could be stereotactically fixed under the two-photon microscope (Figure 6a, b). A recovery period of 2-3 weeks was allowed before *in vivo* microscopic imaging experiments were performed.

**Figure 6:**
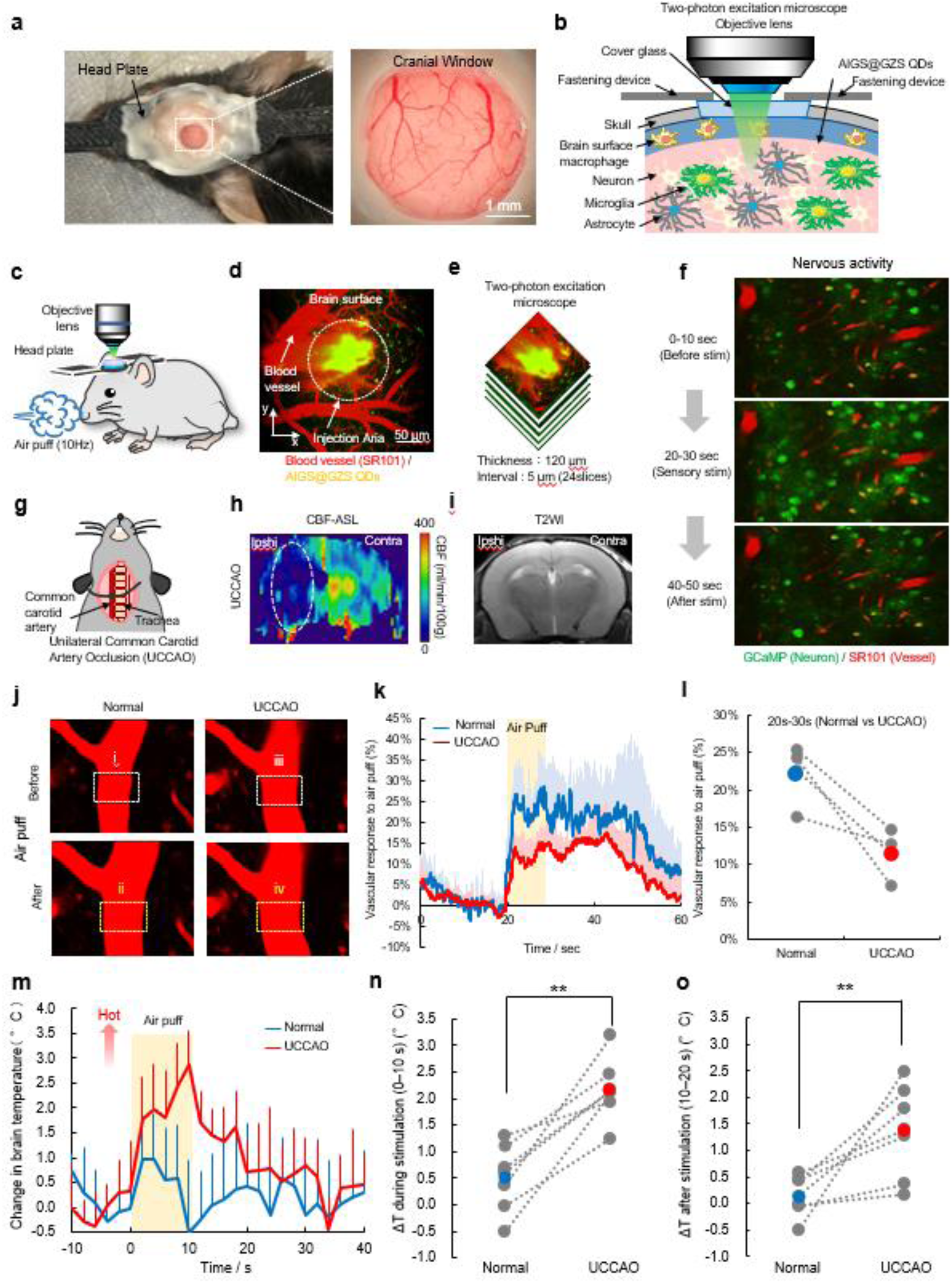
(a) Representative image (left) and enlarged view (right) of a cranial window placed above the somatosensory cortex of a mouse head. Scale bar 1 mm. (b) Schematic diagram of *in vivo* brain temperature measurement using QDs under a two-photon microscope. (c) Schematic diagram of mouse whisker stimulation using an air puff. (d) Representative planar image of the brain surface injected with QDs taken with a two-photon microscope. Scale bar 50 µm. (e) Illustrates the method of imaging QDs using a two-photon microscope. Images were taken over 120 µm in 24 slices at 5 µm intervals from the brain surface. Z stacked average images were obtained using ImageJ. (f) Representative image of neural activity in the barrel cortex when whisker stimulation was applied to a GCaMP transgenic mouse. (g) Schematic diagram of unilateral common carotid artery ligation in a mouse. (h-i) Representative MRI images taken 1 day after unilateral common carotid artery ligation. CBF decreased in the ligated hemisphere. (j) Representative two-photon microscopy images of cerebral blood vessels before and after whisker stimulation under normal conditions and with unilateral common carotid artery ligation. (k) Graph showing the rate of cerebral vasodilation in response to whisker stimulation under normal conditions (blue) and with unilateral common carotid artery ligation (red) (n=4 mice). Data are the average of 4 mice, and error bars indicate standard deviation of the data. (i) Each dashed line during the whisker stimulation period (20-30 seconds after the start of measurement) before and after CCAO indicates data for each individual. (m-o) Temperature measurement results using QDs under normal conditions (blue) and with unilateral common carotid artery ligation (red) (n=6 mice). The horizontal axis is time, and the vertical axis is the rate of temperature change, and the average of 6 mice is shown. Error bars indicate standard deviation. A paired t-test comparing the mean temperature change rates during the stimulation period (20 to 30 seconds after the start of measurement) before and after CCAO (n) and after the stimulation period (30 to 40 seconds after the start of stimulation) (o) showed a significant difference.

Local brain temperature in the barrel cortex was measured before, during and after sensory stimulation using awake mice to remove the effects of anesthesia on neuronal activity and vascular responses (Figure 6c). The plastic bars on the animal’s head were screwed into a custom-made stereotactic apparatus, and the animal was placed on a rotatable styrofoam ball that allowed it to exercise freely while its head was fixed. QD temperature measurement was performed using 4D XYZ-T images. The XYZ-T images had a resolution of 1024 × 1024 pixels and a FOV of 500 μm × 500 μm over depths of 0–120 μm in steps of 5 μm (Figure 6d, e). One volume was acquired every 2 sec, making a total image acquisition time of 60 sec. The image had sufficient depth to cover layers 2 and 3 in the barrel cortex.

In the analysis of the XYZ-T images, the average-intensity projection was calculated in the depth (Z) direction using NIH ImageJ to produce XY-T images. Regions of interest (ROIs) were manually placed on the QDs in the image. The extracted signal was the average PL intensity, either red or green, of all pixels belonging to the ROI. Percent changes in the red-to-green ratio induced by sensory stimulation were normalized with respect to the baseline, which was the average intensity in the 20 sec prior to stimulation. Red-to-green ratio data were averaged over 10 trials for each FOV.

### 2.12. Sensory nerve stimulation and activity measurement

Sensory stimulation was applied to awake mice to induce neuronal activity and perturb the temperature of the barrel cortex. An air puff was delivered to the whiskers at a pressure of 10 psi using an air compressor (NUP-2, AS ONE, Osaka, Japan). Stimulation was performed with rectangular pulses at 10 Hz (50 ms pulse width and 100 ms onset-to-onset interval) generated by a Master-8 (AMPI, Jerusalem, Israel). Each trial was 60 s long (20 s before stimulation, 10 s during the air-puff stimulation, and 30 s after stimulation). Two-photon calcium imaging was performed on the GCaMP6s expressing mice to visualize neuronal activity (Figure 6f). The XY-T calcium images had a resolution of 1024 × 1024 pixels and FOV of 500 μm × 500 μm and were acquired at a frame rate of 2 Hz.

### 2.13. Creation of chronic hypoperfusion model mice

C57BL/6J mice (n=5) were provided with a mixture of air, oxygen, and isoflurane (3-5% for induction and 2% for surgery) via a facemask for unilateral common carotid artery occlusion (UCCAO). After a midline cervical incision, the left common carotid artery (ipsilateral to the cranial window) was isolated from the adjacent vagus nerve and double-ligated with 4–0 silk sutures (Figure 6g).

The reduction in CBF one day after UCCAO was measured using magnetic resonance imaging (MRI). The MRI measurements were performed on mice under isoflurane anesthesia (3-5% for induction and 1.5-2% for measurements) using a 7-Tesla animal MRI system (Bruker, Billerica, Massachusetts, United States). Rectal temperature was continuously monitored and maintained at 37.0 ± 0.5 °C using a heating pad. T2-weighted and CBF-ASL images were acquired, and the CBF-ASL images were analyzed with NIH ImageJ (Figure 6h, i).

### 2.14. Blood flow measurement

To visualize the cerebral blood vessels, 10 mM sulforhodamine 101 (SR101; SIGMA-Aldrich, St. Louis, MO) dissolved in saline was injected intraperitoneally (100 µL) just before the *in vivo* imaging experiments. XY-T blood vessel images had a resolution of 1024 × 1024 pixels and FOV of 500 μm × 500 μm and were acquired at a frame rate of 7 Hz (Figure 6j). The images were acquired at a depth of approximately 100-120 μm (layers 2/3) from the brain surface in the barrel cortex. ROIs were manually placed on each arteriole in the XY-T image with NIH ImageJ. CBF during the air-puff stimulation was evaluated based on the percentage change in the average fluorescence intensity of all pixels belonging to the ROI, and normalized with respect to the baseline (i.e., the average intensity in the 20 sec prior to stimulation). The percentage changes in the CBF were averaged over 10 trials for each animal, and then over all six animals to obtain the mean vascular response.

## Results

### 3.1. Optical properties of AIGS@GZS QDs

To be useful as *in vivo* biosensors, it is necessary that the QDs are stable in aqueous solutions. Since ZnS has a relatively high stability in water, coating QDs with a ZnS shell has been widely employed for QD-based biosensors, such as CdSe@ZnS^25–27^, InP@ZnS^25,26,28^ and Zn-Ag-In-S@ZnS (ZAIS@ZnS)^24,29,30^.

We previously reported^22^that Ag-deficient AIGS QDs exhibited PL spectra consisting of two PL peaks: a broad PL peak originating from defect-site emission and a narrow band-edge emission peak at a shorter wavelength adjacent to the broad peak. This unique PL behavior suggested that a temperature measurement system based on single, low-toxicity, multinary QDs might be developed. However, coating AIGS QD cores with a ZnS layer produced AIGS@ZnS QDs that exhibited only a single broad emission peak (Figure S1) because of additional defect sites at the interface between the AIGS core and ZnS shell^31^. Simlarly while GaSₓ coating enhanced the intensity of band-edge PL, subsequent surface modification of AIGS@GaSₓ QDs with MPA significantly altered the PL spectra, resulting in the complete disappearance of the narrow band-edge emission peak and the emergence of a broad defect-site emission (Figure S2).

Thus, to prepare QDs that have two PL peaks even in aqueous solution, the AIGS QD cores were coated with a Ga–Zn–S shell. As shown in TEM images (Figure 1b and 1f left), the resulting Ga–Zn–S-coated QDs were spherical, with the average diameter increasing from 4.3 nm for the uncoated AIGS QDs to 5.4 nm for the core-shell-structured AIGS@GZS QDs. The uncoated AIGS QDs had a structureless absorption spectrum with an onset at 580 nm (Figure 1c), while the PL spectrum consisted of a sharp emission peak at 560 nm and a broad emission peak around 680 nm (Figure 1d). The former peak was attributed to electron-hole recombination between the conduction band minimum and the valence band maximum (i.e. band-edge emission), and the latter peak originated from the recombination of photogenerated carriers trapped at defect sites (Figure 1e). Surface-coating of the AIGS QDs with a Ga–Zn–S shell layer did not significantly alter the absorption spectrum (Figure 1g), indicating that the energy gap (*E*_g_), size and composition of the AIGS QDs cores were not affected by the shell deposition. In contrast, the PL quantum yield (QY) increased from 12% for uncoated AIGS QDs to 29% for AIGS@GZS QDs, indicating that the photogenerated electrons and holes were confined to the core by the larger energy barrier provided by the Ga–Zn–S shell, that is, the formation of a type-I heterostructure. The wavelength of band-edge emission peak remained nearly constant, appearing at around 570-580 nm for the AIGS@GZS QDs, but the defect-site emission peak was blue-shifted from around 680 nm to around 610 nm, and the relative intensity of defect-site emission to band-edge emission increased after surface coating with Ga–Zn–S (Figure 1h). As the PL properties of QDs are sensitive to the surface conditions, this result suggests that additional defect sites were formed at the AIGS core/Ga–Zn–S shell interface.

Surface modification with MPA enabled uniform dispersion of AIGS@GZS QDs in aqueous solution. As shown in Figure 1g, the absorption spectra of AIGS@GZS QDs remained unchanged before and after the MPA modification. However, MPA modification altered the PL spectrum as the defect-site emission peak was red-shifted from about 610 nm to 640 nm, and the intensity of the band-edge emission peak at around 580 nm decreased (Figure 1h). TEM analysis revealed a slight reduction in the average particle diameter from 5.4 nm to 5.1 nm (Figure 1f, right), while elemental analysis (Table S1) indicated a decrease of the Ga content after dissolving AIGS@GZS QDs in water. The changes observed in the PL spectrum suggest that Ga^3+^ ions contained in the Ga–Zn–S shell were partially leached out during ligand-exchange from OLA to MPA, leading to the observed changes in PL properties. As shown in Figure 1i, MPA-modified AIGS@GZS QDs exhibited yellowish-orange PL under UV light irradiation.

### 3.2. Ratiometric PL sensing of temperature using MPA-modified AIGS@GZS QDs

Figure 1j shows the PL spectra of MPA-modified AIGS@GZS QDs in water at various temperatures. The PL intensity decreased as the solution temperature increased from 5 °C to 45 °C, and the PL QY continuously declined from 22% to 11% (Figure 1m). Figure 1k presents the temperature-dependent changes in the intensities of the band-edge and defect-site emission peaks. Both peaks exhibited a linear decrease in intensity, although the slope was smaller for band-edge emission than for defect-site emission. Although both emission peaks originated from the same QDs, the results indicate that the temperature dependence of the PL intensity differed for each peak due to differences in the emission mechanisms. To further analyze this behavior, we calculated the ratio of the defect-site PL intensity at 640 nm to the band-edge PL intensity at 580 nm, denoted as the red-to-green PL ratio, as a function of solution temperature. As shown in Figure 1l, the red-to-green PL ratio was highly sensitive and decreased approximately linearly from 2.00 to 1.64 with increasing temperature.

### 3.3. Subcutaneous temperature measurement in mouse using AIGS@GZS QDs and in vivo imaging

To evaluate the temperature responsiveness of AIGS@*GZS* QDs, PL imaging was performed while modulating the temperature of a microtube containing AIGS@GZS QDs and a mouse injected subcutaneously with AIGS@GZS QDs (Figure 2a-f). In both cases, the PL intensity decreased as the temperature increased. The PL intensity ratio calculated as the intensity of the edge emission band (550–590 nm) relative to the intensity of the defect-site emission band (690–730 nm), was confirmed to exhibit a high linear correlation with temperature for both experiments (Figure 2e, g).

Subsequently, to evaluate the *in vivo* temperature sensing ability of the AIGS@GZS QDs, AIGS@GZS QDs were injected subcutaneously into the abdomen of a live mouse, and PL imaging was performed at room temperature after either cooling or heating the animal (Figure 3a, b). The temperature was determined by comparing the PL intensity ratio, derived from the PL images to the calibration curve in Figure 3c. The temperature estimated using the AIGS@GZS QDs agreed well with the temperature measured using an infrared camera. These results indicate that the PL intensity ratio of AIGS@GZS QDs provides accurate measurements of *in vivo* temperature.

### 3.4. NIR laser excitation of AIGS@GZS QDs and temperature measurement

Due to their small size, AIGS@GZS QDs have the potential to provide high spatial resolution imaging of *in vivo* temperature. Many fluorescent probes have been shown to undergo multiphoton excitation under NIR laser illumination. NIR light can penetrate tissue to a depth of around 1mm, so it is widely used for high-resolution *in vivo* imaging^33,34^. To characterize the excitability of AIGS@GZS QDs under NIR laser excitation, we performed *in vitro* measurements.

It was confirmed that the AIGS@GZS QDs were sufficiently excited over a broad range of the NIR spectrum (700–1040 nm) (Figure 4a–c). While the excitation efficacy peaked at around 740–810 nm, the AIGS@GZS QDs were also sufficiently excited in the range 900–1000 nm. The latter wavelength range is known to exhibit low absorption in brain tissue and thus is less harmful for continuous *in vivo* time-lapse imaging. Also, since cost-effective NIR lasers with a fixed 920 nm wavelength are commercially available, we selected 920 nm as the target wavelength to further evaluate the proposed temperature imaging technique.

### 3.5. Two-photon PL characteristics and temperature measurement

To evaluate whether AIGS@GZS QDs could be used for temperature measurement with a two-photon excitation laser microscope, a 1.5% agarose gel QD phantom was constructed and analyzed. To directly compare temperature data obtained with QDs and two-photon microscopy to conventional temperature measurement techniques, a thermocouple probe was inserted into the phantom. The phantom with the thermocouple was placed on a temperature control device, and the PL intensity of the QDs was captured at temperatures ranging from 25°C to 40.1°C using a two-photon excitation laser microscope (Figure S3a). A scatter plot of the red-to-green PL intensity ratio against the thermocouple measurements is presented in Figure S3b. The two data sets were linearly correlated with R^2^=0.9896. This result directly demonstrated that the QDs function effectively as temperature sensors under two-photon excitation.

An additional experiment was conducted to assess the temporal sensitivity of the QD-based temperature measurement system. The QD phantom was placed on a temperature control device, and the thermocouple was used to monitor the temperature as it fluctuated between 29.9°C and 30.1°C, repeating the cycle three times over 60 seconds. Changes to the PL intensity of the QDs were measured using a two-photon microscope, and the PL intensity ratio (red-to-green) was calculated. The temperature data obtained from the QDs closely followed the thermocouple-measured temperature fluctuations. This finding confirmed that the QDs were capable of tracking temperature fluctuations on the scale of seconds (Figure S3c).

Next, to evaluate whether AIGS@GZS QDs could be used for brain temperature measurement with a two-photon excitation laser microscope, the QDs were injected into mouse brain. To directly compare temperature data obtained with QDs and two-photon microscopy to conventional temperature measurement techniques, a thermocouple probe was also inserted into the mouse brain. The mouse was placed on a temperature control device, and the PL intensity of the QDs was captured at temperatures ranging from 24.0°C to 35.1°C using a two-photon excitation laser microscope (Figure 4d). A scatter plot of the red-to-green PL intensity ratio against the thermocouple measurements is presented in Figure 4e. The two data sets were linearly correlated with R^2^=0.8649. This result directly demonstrated that the QDs function effectively as brain temperature sensors under two-photon excitation.

### 3.6. Observation after intracerebral injection of AIGS@GZS QDs in mice

To investigate the distribution of QDs locally injected into the brain via a fine glass needle, immunofluorescence-stained tissue sections were analyzed. A schematic diagram of the experimental method is shown in Figure 5a. Brain slices were stained for GFAP (astrocytes), Iba1 (microglia), and tubulin β3 (neurons). Representative images acquired using a confocal microscope with a 10× objective lens are shown in Figure 5b. The QDs were observed near the site of injection. Imaging at 20× magnification revealed no significant morphological differences in astrocytes, microglia, or neurons between the QD-injected and saline-injected conditions (Figure 5c). QDs were not observed inside cells; instead, they remained in the extracellular space of the brain parenchyma for three days post-injection.

### 3.7. Temperature measurement in the brain parenchyma of normal mice after sensory nerve stimulation using AIGS@GZS QDs

A cranial window was implanted in the head of wild-type and GCaMP6s mice (Figure 6a). A schematic diagram of the imaging method using a two-photon excitation laser microscope in the awake state is shown in Figure 6b. Measurements using the two-photon excitation laser microscope were conducted in the barrel field of the somatosensory cortex. To evoke neural activity in the barrel field, air-puff stimulation was applied to the whiskers (Figure 6c). Representative images of neural activity in GCaMP6s mice are shown in Fig. 6f. Changes to the fluorescence intensity from individual neurons indicated an increase in neuronal activity upon stimulation.

Next, the vasodilation response of arterioles in the barrel field during sensory stimulation was examined. Figure 6j shows representative blood vessel images and regions of interest (ROIs) before and after air-puff stimulation in normal mice. In normal mice, an increase in blood vessel diameter was observed following sensory input. Similarly, the mean fluorescence intensity of the blood vessels showed a rapid increase immediately after sensory input, with a maximum response of approximately 28% (Figure 6k). Many previous studies have demonstrated that cerebral blood vessels dilate in response to neural activity, and our findings support those prior reports.

To investigate the relationship between neural activity and brain temperature, we conducted temperature measurements in six mice using QDs. A representative two-photon excitation laser microscope image of QDs near the brain surface is shown in Figure 6d. Based on the acquired images, the red-to-green PL intensity ratio was calculated and converted into brain temperature changes. The average data from six animals is shown in Figure 6m. Brain temperature gradually increased after whisker stimulation, reaching a peak increase of 1.0℃ in 2 seconds. The temperature then gradually decreased, returning to the baseline level in approximately 30 seconds after the start of measurement.

### 3.8. Temperature measurement in the brain of chronic hypoperfusion model mice during sensory stimulation

Next, we conducted brain temperature measurements during sensory stimulation in chronic hypoperfusion model mice. It has been reported that cerebral blood vessels dilate after common carotid artery occlusion (CCAO), and cerebral vascular reactivity to CO₂ inhalation and neural activity is diminished in this model^34,35^. It has also been reported that there is no significant neuronal loss for at least one week after CCAO^36^. Thus, comparing chronic hypoperfusion model mice with normal mice allows for an analysis of temperature changes during neural activation under conditions of normal and impaired cerebrovascular reactivity.

First, to confirm whether the decrease in vascular reactivity observed in previous studies was also present in our model, we measured changes in blood vessel fluorescence intensity using two-photon microscopy. For vascular imaging, sulforhodamine 101 was intravenously injected to fluorescently label the vascular lumen, and two-photon imaging was performed before (normal state) and one day after CCAO. The same measurement region and identical blood vessels were targeted before and after CCAO. Representative images of brain blood vessels before and after sensory stimulation, along with ROIs, are shown in Fig. 6j. In the normal state before CCAO, a comparison between ROI 1 and ROI 2 showed that the blood vessels dilated after stimulation. Comparing the resting-state blood vessels at ROI 1 and ROI 3 revealed that the vessel diameter increased after CCAO. A comparison of pre- and post-stimulation vessel diameters showed that blood vessel dilation was more pronounced before CCAO (ROI 1 vs. ROI 2) than after CCAO (ROI 3 vs. ROI 4). This suggests that after CCAO, resting-state blood vessels were wider, but the dilation response to neural activation was diminished.

The fluorescence intensity data from four mice were averaged, and the fluorescence intensity ratio was standardized to the mean intensity during the 20 seconds before stimulation. The fluorescence intensity ratio during sensory stimulation was higher before CCAO than after CCAO (Figure 6k). A paired t-test comparing the mean fluorescence intensity ratio during stimulation before and after CCAO found a significant difference between the control and hypoperfused groups (Figure 6l). These results are consistent with previous reports indicating that cerebrovascular reactivity is reduced in chronic hypoperfusion model mice during neural activation.

After confirming the effectiveness of the chronic hypoperfusion mouse model as an experimental intervention, we performed QD temperature measurements in the same region before and after CCAO. The temperature was determined from the red-to-green PL intensity ratio of the QDs. Based on the calibration obtained from the ex vivo brain phantom experiment (Figure 4e), the PL intensity ratio was converted into temperature. Under normal conditions (before CCAO), the average temperature increase during the stimulation period (0–10 s after stimulation onset) was 0.51°C. In contrast, after CCAO, the average temperature increase during the same period reached 2.2°C, representing a more than fourfold increase compared with the normal condition (Figure 6m). A paired t-test comparing the average red-to-green PL intensity during the stimulation period (20–30 seconds after measurement onset) before and after CCAO showed a significant difference (Figure 6n). Furthermore, even after the stimulation period (30–40 seconds after measurement onset), a significant temperature increase was sustained in the CCAO condition (Figure 6o). This raises the possibility of directly analyzing brain thermal dynamics at the microregional level, which up till now has been discussed primarily through theoretical or indirect evidence.

## Discussion

In this study, we successfully developed water-soluble, low-toxicity QDs that exhibit both band-edge (green) and defect-state (red) emissions, for which the red-to-green emission peak intensity ratio changes linearly with ambient temperature. Conventional QD-based temperature sensors relied on correlations between PL intensity and temperature^37,38^, but those approaches were highly susceptible to tissue-dependent absorption and scattering effects, making accurate temperature measurements difficult. Methods based on detecting slight temperature-dependent shifts in the PL peak wavelength have also been proposed, but such approaches are sensitive to detector resolution limits and noise, making highly accurate temperature measurements challenging^4,6,7^. In contrast, the method developed in this study is based on the linear correlation between the red-to-green PL intensity ratio and temperature rather than the absolute intensity or wavelength of the PL peak. By calculating the red-to-green PL intensity ratio as an integrated intensity ratio, instability caused by fluctuations in the properties of the QDs generated during the synthesis process are substantially reduced. This strategy minimizes tissue-derived optical artifacts to enable stable and highly accurate temperature measurements in vivo.

Previously, in vivo temperature measurements were feasible using biological imaging systems such as IVIS, and useful results were obtained through comparisons with thermography. In the future, red-shifting of QDs may enable temperature measurements in deeper tissues and organs.

To fully exploit the advantages of QDs, we combined them with two-photon excitation microscopy and applied them to brain temperature measurements. In vivo imaging using two-photon microscopy offers high spatiotemporal resolution and enables repeated three-dimensional measurements. As such temperature measurements at a spatial resolution of 1 μm × 1 μm became possible with a temporal resolution on the order of several tens of milliseconds. Furthermore, repeated volumetric temperature measurements from the brain surface to a depth of 120 μm were possible at 2-second intervals. Existing thermocouples provide rapid response times but suffer from poor spatial resolution^39–41^. In addition, MRI and nanodiamond-based methods applicable to in vivo imaging have temporal resolutions on the order of seconds, limiting their ability to track rapid temperature fluctuations^41–43^. In contrast, QDs with high spatiotemporal resolution two-photon microscopy represent an extremely useful tool for tracking temperature dynamics within local brain circuits.

The developed technology was applied to measure temperature changes in active brain regions. To achieve this, it was important to position QDs within the target perivascular regions. This was accomplished using COOH surface-modified QDs (ref: Yukawa et al.). Immunostaining confirmed that the quantum dots were not taken up into brain cells. Furthermore, observation of vascular structures using two-photon microscopy confirmed that the quantum dots were localized within the intended perivascular regions. Stable retention within tissues for more than one week was also confirmed (Data not shown).

To evaluate brain temperature dynamics under physiological conditions, we applied measurement system to awake mice. Anesthesia is frequently used in conventional rodent experiments, but the anesthetic itself can affect brain temperature measurements^45–47^. In contrast, the awake mouse model used in this study did not require heating devices and enabled the measurement of brain temperature fluctuations under conditions closer to natural physiological states.

Measurements under physiological conditions are important for understanding mechanisms of heat production and heat dissipation in the brain. The brain is a highly metabolically active organ that generates large amounts of heat^10,11,17,18^. In contrast, cerebral blood vessels are known to be cooler than the brain parenchyma and function to absorb heat from surrounding tissues and dissipate it outside the body, even under resting conditions^11,14,17^.

Furthermore, an important physiological phenomenon unique to the brain is neurovascular coupling (NVC), which is a local increase in cerebral blood flow associated with neural activity^19^. Initially, increases in cerebral blood flow induced by NV coupling were thought to compensate for glucose and oxygen consumed during neural activity^48,49^. However, subsequent research studies revealed that neural activity induces cerebral blood flow increases far exceeding actual metabolic demand^50,51^, and the physiological significance of this excessive blood flow response has long been debated^19,48,49^. It has been suggested that the blood flow response is involvement in brain is involved in the removal of heat generated during neural activity^11,14,17,52^.

Because NV coupling occurs at the micrometer-scale, direct biological verification of its physiological significance requires in vivo temperature measurement technologies with high spatiotemporal resolution. Unfortunately, conventional MRI and thermocouples have limited in spatiotemporal resolution, making detailed evaluation of local and transient temperature changes difficult^53^. By utilizing the linear correlation between the red-to-green PL intensity ratio and temperature, the QD-based method proposed in this study greatly reduced the optical noise due to dural thickening beneath cranial windows, thereby enabling in vivo brain temperature measurements at the micrometer scale.

In this study, we observed an average brain temperature increase of approximately 0.51°C in the somatosensory cortex during sensory stimulation. This result is consistent with temperature increases during brain activation reported in human MRI studies and animal thermocouple experiments^17,52,54^. For example, previous mouse sensory stimulation experiments reported temperature increases of approximately 0.35°C^52^. These findings indicate that the QD-based temperature measurement method developed in this study successfully captured local heat production associated with neural activity at high spatiotemporal resolution.

Temperature sensing using QDs allows the temperature to be measured at a spatial resolution of 1 μm × 1 μm, greatly surpassing the resolution of conventional thermocouple probes which typically have diameters reaching several hundred micrometers, as well as MRI, which remains limited to millimeter-scale resolution at best. Furthermore, two-photon microscopy enables three-dimensional temperature mapping within living tissues. Owing to the high spatiotemporal resolution of the method, local temperature elevations can be detected with high precision, which may explain why the temperature increases observed in this study were large greater than those reported in previous thermocouple-based studies with thermocouples or MRI, local temperature hotspots are averaged together with surrounding tissues where there is no heating, whereas the present method may have captured local heat production more directly.

In chronic hypoperfusion mice, the average temperature increase during neural activity was elevated in comparison to healthy control mice, and transient peaks exceeding 2–3°C were observed in some cases. These findings represent the first direct in vivo demonstration, at the microregional level, that neural activity–associated increases in cerebral blood flow function not only to supply oxygen and glucose, but also as a physiological mechanism for dissipating locally generated heat. By demonstrating markedly amplified neural activity–dependent temperature elevations under conditions of impaired vascular reactivity, the present study directly verified that increased cerebral blood flow functions as a thermoregulatory mechanism for brain. In other words, cerebral blood vessels may function not only as pathways for the delivery and clearance of substances associated with neural activity, but also as an extremely efficient “brain temperature control system” that dynamically modulates local cooling functions in response to changes in heat production (Figure 7).

**Figure 7:**
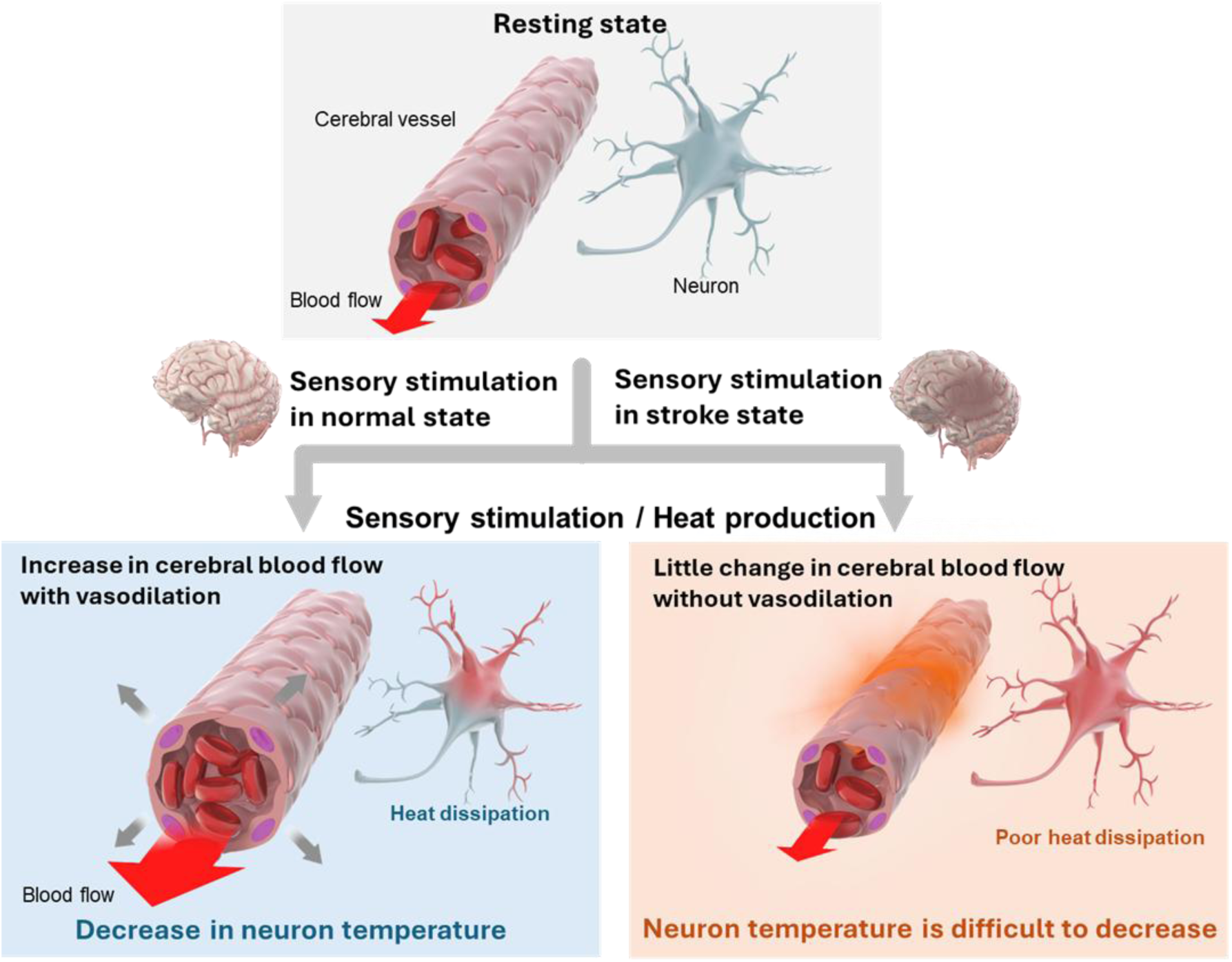
Schematic diagram of heat production due to neural activity and the mechanism by which brain temperature is regulated by cerebral blood flow. Under normal conditions, the rise in brain temperature caused by heat production due to neural activity is dissipated by an increase in cerebral blood flow due to vasodilation. However, during a stroke, the rate of vasodilation decreases, so the increase in cerebral blood flow is smaller, heat is not dissipated, and brain temperature is less likely to decrease.

Consistent with the excessive temperature elevations observed in the chronic hypoperfusion model in the present study, some clinical studies have also reported elevated brain temperatures during the acute phase of stroke^49,55,56,57^. Even a 1°C increase in brain temperature under stroke conditions is known to accelerate infarct expansion38, suggesting that transient and localized temperature elevations may have a profound effect on neurons and glial cells. These findings imply that localized “thermal runaway” occurring during cerebrovascular dysfunction may indicate a previously unrecognized pathological mechanism underlying the progression of brain injury. Furthermore, the microscale brain temperature measurement technology developed in this study is expected to contribute not only to stroke research, but also to the understanding of disease mechanisms and the development of therapeutic strategies for various disorders associated with cerebrovascular dysfunction, including dementia and other neurodegenerative diseases.

Another important advantage of the QD-based temperature measurement method developed in this study is its compatibility with simultaneous imaging of other fluorescent dyes and fluorescent proteins under two-photon microscopy. This enables simultaneous measurements of calcium responses, vascular dilation, and temperature changes within the same FOV during neural activity. In particular, the method is expected to serve as a highly useful platform for understanding the interactions between neural activity, cerebral blood flow, and heat production.

In the future, the method may also enable real-time visualization of how heat generated by neural activity diffuses into surrounding blood vessels and is dissipated. This raises the possibility of directly analyzing brain thermal dynamics at the microregional level, which up till now has been discussed primarily through theoretical or indirect evidence. In particular, the cerebral blood flow–mediated heat dissipation mechanism demonstrated in this study may eventually be visualized directly with high spatiotemporal resolution, potentially leading to a new physiological understanding of the relationship between neural activity and cerebral blood flow.

In addition, the method may be applicable to a variety of brain cell types, including microglia and tumor cells. Less invasive delivery of QDs using intravenous administration or blood–brain barrier penetration technologies may become feasible in the future. By combining technologies for efficient QD delivery into neurons and glial cells, direct measurements of intracellular temperature dynamics may become possible, opening broad applications in neuroscience and brain disease research.

## Conclusion

In this study, we presented a QD-based temperature sensing system capable of microscale brain temperature measurements in vivo. By combining QDs with two-photon microscopy, we achieved simultaneous measurements of brain temperature, neural activity, and cerebrovascular response in awake mice, and successfully visualized local heat production associated with neural activity at high spatiotemporal resolution.

Furthermore, using a chronic hypoperfusion model with impaired cerebrovascular reactivity, we demonstrated excessive neural activity–dependent temperature elevations and directly showed in vivo that increased cerebral blood flow functions as a thermoregulatory mechanism that dissipates local heat. The findings support the existence of a “heat dissipation mechanism” as a previously unrecognized physiological effect of NV coupling and suggest that cerebral blood vessels may function as an extremely efficient brain temperature control system.

The microscale brain temperature measurement technology developed in this study is expected to provide new insights into pathological conditions associated with cerebrovascular dysfunction, including stroke, dementia, and neurodegenerative diseases, and may also contribute to the development of therapeutic strategies focused on intracerebral thermal dynamics.

