## Supporting Information for "*In vivo* brain temperature measurement using quantum dot imaging temperature sensors"

### Abstract

Temperature regulation in the brain is essential for maintaining neuronal function and preventing thermally induced damage. Here, we report the development and *in vivo* application of quantum dots (QDs) to high-resolution thermometry in the mouse brain using two-photon excitation microscopy. These QDs, via the red-to-green photoluminescence (PL) intensity ratios, enabled stable temperature measurements in both normal and chronically hypo-perfused cerebral tissue. Our findings show that localized neuronal activity leads to transient heat generation, which is rapidly dissipated by cerebrovascular responses. In a chronic hypoperfusion model, impaired vascular function resulted in exaggerated and prolonged brain temperature elevations. This thermometry system provides unprecedented insight into the mechanisms of cerebral thermoregulation and highlights the importance of vascular cooling in protecting the brain from heat-induced stress, particularly in pathological conditions such as stroke.

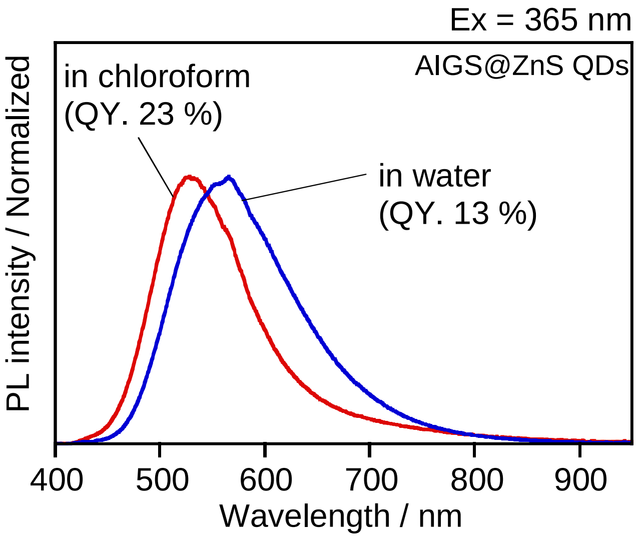


**Figure S1.** PL spectra of AIGS@ZnS QDs. The QDs modified with OLA and MPA were uniformly dispersed in chloroform and water, respectively.

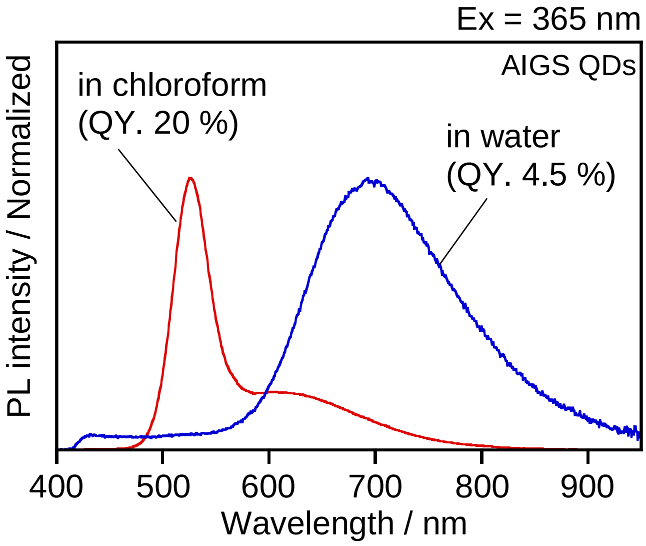


**Figure S2.** PL spectra of AIGS and AIGS@$GaS_x$ QDs. The QDs modified with OLA and MPA were uniformly dispersed in chloroform and water, respectively.

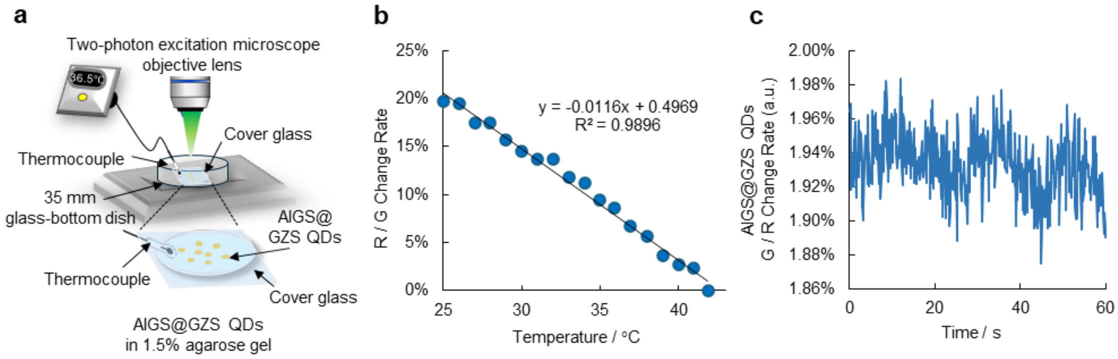


**Figure S3.** (a) Schematic diagram of QD temperature calibration using a gel phantom with QDs embedded in 1.5% agarose gel. (b) Graph comparing the measured value by a thermocouple probe inserted into the gel with the rate of change in the QD PL intensity ratio. $R^2 = 0.9896$. (c) Rate of change in the QD PL brightness ratio with respect to fluctuations in heater temperature. The heater temperature ranged from 29.9℃ to 31.1℃.

**Table S1.** Chemical composition of AIGS@GZS QDs surface-modified with OLA and MPA.

| ligand | Ag | In | Ga | S | Zn | Ga/Ag |
|---|---|---|---|---|---|---|
| OLA | 5.5 | 4.1 | 11 | 51 | 28 | 1.9 |
| MPA | 5.0 | 6.2 | 6.7 | 54 | 28 | 1.3 |
